# Doxycycline inhibits α-synuclein-associated pathologies *in vitro* and *in vivo*

**DOI:** 10.1101/2020.11.06.371229

**Authors:** Antonio Dominguez-Meijide, Valeria Parrales, Eftychia Vasili, Florencia González-Lizárraga, Annekatrin König, Diana F. Lázaro, Annie Lannuzel, Stéphane Haik, Elaine Del Bel, Rosana Chehín, Rita Raisman-Vozari, Patrick P Michel, Nicolas Bizat, Tiago Fleming Outeiro

**Author notes:** These authors contributed equally to this work (Antonio Dominguez-Meijide, Valeria Parrales). Co-senior authors, **Email:** Prof. Dr. Tiago F. Outeiro, Department of Experimental Neurodegeneration, University Medical Center Goettingen, 37073 Goettingen, Germany, or Dr. Nicolas Bizat, Institut du Cerveau et de la Moelle, Hôpital de la Pitié-Saplêtrière, 47, Bd de l’Hôpital, Paris F-75013, France.

## Abstract

Parkinson’s disease (PD) and dementia with Lewy bodies (DLB) are neurodegenerative disorders characterized by the misfolding and aggregation of alpha-synuclein (aSyn). Doxycycline, a tetracyclic antibiotic shows neuroprotective effects, initially proposed to be due to its anti-inflammatory properties. More recently, an additional mechanism by which doxycycline may exert its neuroprotective effects has been proposed as it has been shown that it inhibits amyloid aggregation. Here, we studied the effects of doxycycline on aSyn aggregation *in vivo, in vitro* and in a cell free system using real-time quaking induced conversion (RT-QuiC). Our results show that doxycycline decreases the number and size of aSyn aggregates in cells. In addition, doxycycline inhibits the aggregation and seeding of recombinant aSyn, and attenuates the production of mitochondrial-derived reactive oxygen species. Finally, we found doxycycline induces a cellular redistribution of the aggregates in an animal model of PD that is associated with a recovery of dopaminergic function. In summary, we provide strong evidence that doxycycline treatment may be an effective strategy against synucleinopathies.

## Introduction

Parkinson’s disease (PD) is the second most common neurodegenerative disease, affecting more than ten million people worldwide (Tysnes & Storstein, 2017). PD and some other synucleinopathies like dementia with Lewy bodies (DLB) are characterized by the presence of intraneuronal aggregates composed of the protein alpha-synuclein (aSyn; (Spillantini et al, 1998)) and the overexpression of this protein in neurons contributes to these pathologies (Mehra et al, 2019; Outeiro et al, 2019).

Monomeric aSyn is an intrinsically disordered neuronal protein highly enriched in presynaptic nerve terminals supporting the hypothesis that aSyn is a regulator of dopaminergic (DA) neurotransmission (Abeliovich et al, 2000). aSyn deficient mice exhibit a reduction in striatal dopamine and an attenuation of dopamine-dependent locomotor response. The molecular mechanisms involved in synucleinopathies, from aSyn misfolding and aggregation to the various cellular effects and pathologies associated remains unclear (Brás et al, 2020). This may happen most likely due to aggregated aSyn disrupting vesicular transport, impairing autophagy, causing mitochondrial deficits and affecting vesicle recycling, neuronal plasticity and synaptic integrity (Bonini & Giasson, 2005; Liang & Tamm, 2018; Lindström et al, 2017; Murphy et al, 2000; Winslow et al, 2010). Furthermore, aggregate forms of aSyn can self-propagate within neurons and to interconnected neural networks throughout the nervous system in a prion-like manner resulting in the spread of pathology throughout the brain and the progression of PD (Angot et al, 2012).

Doxycycline is a wide-spectrum antibiotic that belongs to the group of the tetracyclines reported to be neuroprotective (Reglodi et al, 2017; Santa-Cecília et al, 2019). The neuroprotective effect was proposed to be due to its anti-inflammatory properties similar to minocycline, another tetracycline antibiotic (Bortolanza et al, 2018; Cronin & Grealy, 2017). In fact, in animal models of PD, it has been seen that tetracycline derivatives can protect DA neurons in the substantia nigra (Cho et al, 2009; Du et al, 2001), possibly by restraining the glial response in the 6-OHDA animal model of PD (Lazzarini et al, 2013). In addition, doxycycline inhibits fibril formation of amyloid beta (Aβ), prion protein (PrP), β-microglobulin and aSyn (Costa et al, 2011; Giorgetti et al, 2011; González-Lizárraga et al, 2017; Hannaoui et al, 2014; Schmitz et al, 2016). More recently, it has been proposed that doxycycline may also act against amyloid aggregation in neurodegenerative diseases (Socias et al, 2018; Stoilova et al, 2013). Even though the neuroprotective effects of doxycycline are clear in various model systems, the mechanism of its action remains to be elucidated.

The nematode *Caenorhabditis elegans* (*C. elegans*) possess a well-known neuronal system and has emerged as a valuable experimental *in vivo* model for understanding the neurobiological processes involved in neurodegenerative diseases like Parkinson, Alzheimer’s, Huntington’s and prion diseases (Alexander et al, 2014; Bizat et al, 2010; Dexter et al, 2012; Gaeta et al, 2019; Hannan et al, 2016; Rand & Nonet, 1997; White et al, 1986). To study *in vivo* the potential interactions that may exist between doxycycline and aSyn, we developed transgenic models expressing the human wild-type aSyn in the GFP-tagged DA neurons. In the present study, we show that doxycycline reduces the formation of aSyn inclusions by inhibiting the seeding process and by disrupting the seeds, as observed by RT-QuiC and Thioflavin T (ThT) seeding assays. In addition, doxycycline inhibits the production of reactive oxygen species by mitochondria. Importantly, we found that, *in vivo*, doxycycline modifies the cellular distribution of aSyn inclusions in DA neurons and restores altered motor behaviours.

## Results

### Doxycycline reduces aSyn aggregation *in a cell free system*

In order to evaluate the effect of different doxycycline concentrations on aSyn aggregation we used the RT-QuiC aggregation assay (Wilham et al, 2010). The results obtained showed a dose dependent decrease in ThT fluorescence intensity. Basal values decreased with higher concentrations of doxycycline pointing not only to inhibition of aggregation but also to a possible change in the conformation of the aggregated aSyn (Figure 1A).

**Figure 1.**
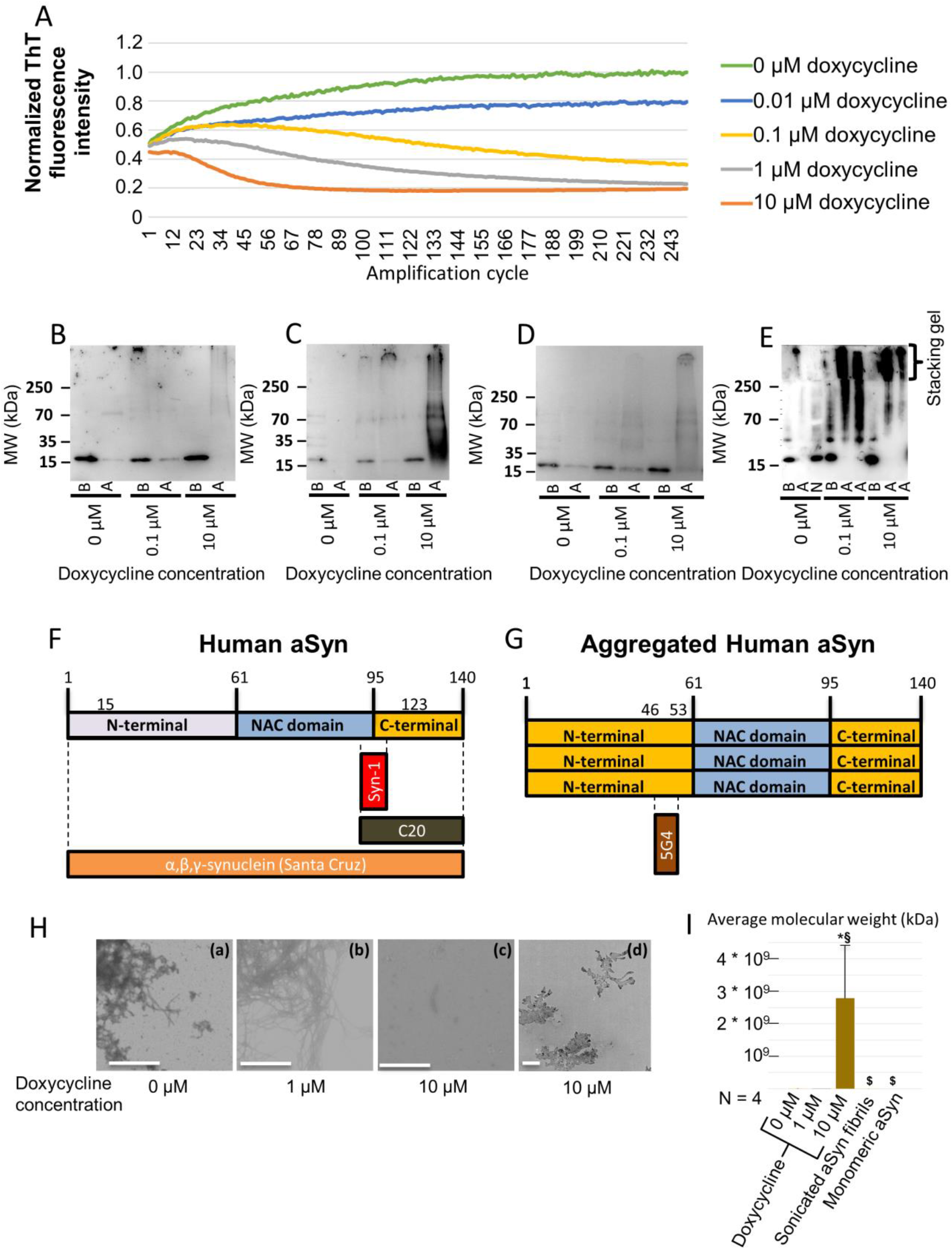
Effects of the presence of different doxycycline concentrations on *in vitro* amplification of aSyn by RT-QuiC. **A**. Effect of doxycycline on aSyn aggregation by RT-QuiC. Increasing concentrations of doxycycline lead to lower ThT fluorescence intensity. **B.** SDS-PAGE membrane of the RT-QuiC products developed with the BD anti-aSyn antibody. (*b: Sample before amplification. a: Sample after amplification*). **C.** SDS-PAGE membrane of the RT-QuiC products developed with the anti-alpha beta gamma aSyn antibody. (b: *Sample before amplification. a: Sample after amplification*). **D.** SDS-Page membrane of the RT-QuiC products developed with the 5g4 anti-aggregated aSyn antibody. (*b: Sample before amplification. a: Sample after amplification*). **E**. Semi-Denaturing gel membrane of the RT-QuiC products. (*b: Sample before amplification. a: Sample after amplification. n: Negative control for RT-QuiC*). Presence of doxycycline during RT-QuiC amplification lead to the formation of higher molecular weight species above equimolar concentrations. **F.** Epitopes for the anti-human aSyn (Syn-1 and C20) and anti-αβγSyn antibodies. **G.** Epitope for the 5G4 anti-aggregated human aSyn antibody. **H.** Electron microscopy pictures of the samples obtained in the RT-QuiC. (a) In samples without docycycline, abundant fibrillar structures are observed. Scale bar 900 nm. (b) Below equimolar concentrations (1 μM) abundant fibrillar structures are still observed. Scale bar 900 nm. (c) Above equimolar concentrations (10 μM) few fibrils are seen. Scale bar 900 nm. (d) Above equimolar concentrations large non-fibrillar structures are observed. Scale bar 900 nm. **I.** Average molecular weight obtained by DLS. The presence of 10 μM doxycycline leads to the formation of higher molecular weight species. Data presented as mean ± SD. *p < 0.05 in comparison with 0 μM Doxycycline. §p< 0.05 in comparison with 1 μM Doxycycline. $p< 0.05 in comparison with 10 μM Doxycycline. Monomers displayed an average value of 16 kDa approximately.

Samples collected before and after the RT-QuiC reaction were run on SDS-PAGE gels and, upon immunoblotting with the Syn-1 antibody, we observed a clear signal corresponding to monomeric aSyn in the samples prior to RT-QuiC amplification (Figure 1B). On the other hand, after RT-QuiC amplification we generally observed no monomeric, neither multimeric aSyn signal. We then incubated the membranes with other anti-aSyn antibodies that bind to larger epitopes(Perrin et al, 2003) (pan αβγ-Synuclein or C20-Synuclein), and detected a smear in samples incubated with 10 μM doxycycline (Figure 1C). As this suggested the presence of aggregated species, we incubated the membranes with the 5G4 anti-aggregated aSyn antibody, observing again a smear in the samples treated with 10 μM doxycycline (Figure 1 D).

Interestingly, when the samples were run in a semi-denaturing polyacrylamide gel electrophoresis (SDD-PAGE) tricine gel, samples amplified in the presence of 0.1 μM doxycycline showed smears ranging from approximately 35 kDa all the way up to the top of the gel while samples amplified in the presence of 10 μM doxycycline showed only higher molecular weight species (Figure 1E). The absence of the smear when we probed with the Syn-1 antibody might be due to the differences in the epitopes recognized, as pan αβγ-Synuclein and C20-Synuclein bind to the C-terminus, and the 5G4 to a different region of the protein (Figure 1F,G). As it has been previously reported that, above equimolar concentrations, doxycycline, may change the conformation of the protein, leading to the formation of non-fibrillary species we then analysed the samples using electron microscopy (EM) (González-Lizárraga et al, 2017). Samples amplified without doxycycline or with 1 μM doxycycline showed the presence of fibrils, while samples amplified in the presence of 10 μM doxycycline showed some fibrils and large non-fibrillary structures. In the RT-QuiC assay, 1 μM doxycycline showed an effect after more than 25 cycles (Figure 1 H).

Next, in order to further assess the size of the aggregated species formed, we used dynamic light scattering (DLS). We detected very high molecular weight (HMW) species in samples amplified in the presence of 10 μM doxycycline (Figure 1I, Supplementary Figure S1). We also assessed the average molecular weight of the monomers used for amplification, obtaining a result of 16.05 ± 1.8 kDa, consistent with the reported molecular weight of 14.5 kDa.

### Doxycycline reduces seeded aSyn aggregation and the levels of reactive oxygen species generated by mitochondria

Next, in order to further investigate the effect of doxycycline on aSyn aggregation, we tested whether it affected the seeded aggregation of monomeric aSyn, as this might relate to the spreading of aSyn pathology as disease progresses. In vitro, in cell free models, the process of aSyn aggregation follows a typical sigmoidal curve, with a latency time of 16 hours followed by an exponential phase that reached a stationary state around 44 hours (Figure 2A), as previously described (Ávila et al, 2014). In contrast, the presence of aSyn aggregates triggered a rapid aggregation of monomeric aSyn, abolishing the lag phase. However, when doxycycline was added to the reaction, the increase in ThT fluorescence intensity was completely blocked. This suggests that doxycycline *“poisons”* the seeding of aSyn aggregates species.

**Figure 2.**
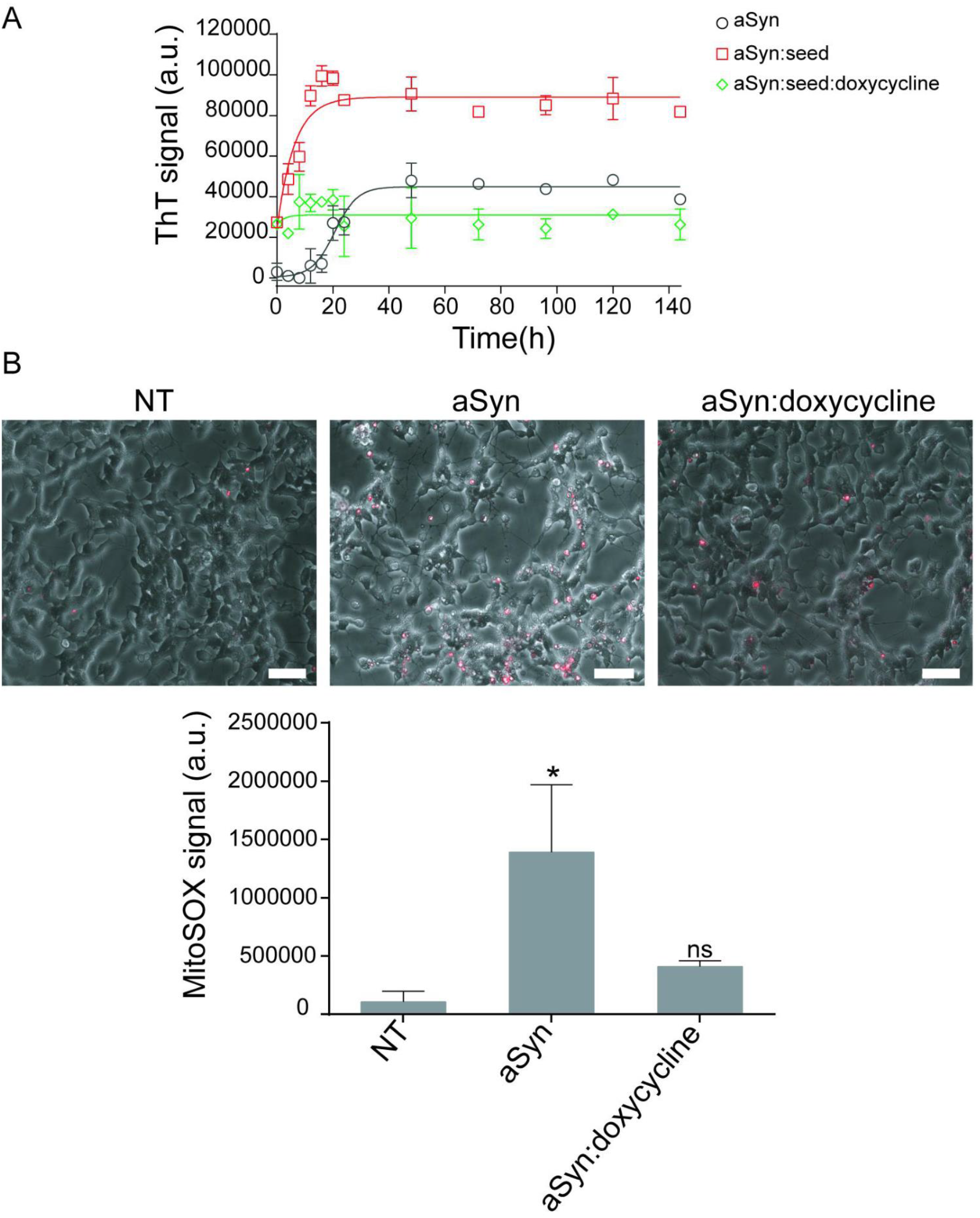
Doxycycline interferes with aSyn seeding and mitochondrial ROS production. **A.** Doxycycline inhibits αSyn seeding *in vitro*. Fresh solutions of monomeric αSyn were seeded with aggregates and incubated in the absence (empty squares) or presence of doxycycline (empty diamonds) at 37 °C under orbital agitation. Aggregation was assayed by ThT fluorescence emission. Unseeded aggregation kinetics are also shown (empty circles). **B.** Intracellular ROS production visualized with MitoSOX Red probe in differentiated SH-SY5Y cells treated with aSyn aggregates produced in the presence or the absence of doxycycline. Scale bar: 100 μm. MitoSOX Red fluorescence intensities in the same culture conditions as in A *p < 0.05 *vs* NT.

Mitochondrial dysfunction leads to the production of reactive oxygen species and can directly interact with variety of specific proteins to be involved in neurodegenerative disorders (Lin & Beal, 2006). For example, in PD aSyn aggregates may induce neurodegeneration by causing a mitochondrial dysfunction and, thereby, the increased production of reactive oxygen species (ROS) (Park et al, 2020; Santos & Outeiro, 2020; Szegő É et al, 2019; Wang et al, 2019). For this reason, we measured intracellular ROS production in differentiated SH-SY5Y cells treated with aggregates formed in the absence or presence of 100 μM doxycycline. SH-SY5Y cells, as most neuronal cell lines, are more susceptible to oxidative stress than glial cells (Wang & Michaelis, 2010). Using MitoSOX Red, we observed that SH-SY5Y cells exhibited a significant increase in intracellular ROS when in the presence of aSyn aggregates (Figure 2B). However, when the cells were treated with aggregated aSyn, formed in the presence of doxycycline, this effect was almost completely blocked.

### Doxycycline decreases the number of aSyn inclusions in cells

In order to assess the effect of doxycycline treatment on aSyn aggregation in cell cultures, we used the SynT/Synphilin-1 model of inclusion formation (Lázaro et al, 2014; Masaracchia et al, 2020). SynT is a C-terminally modified variant of aSyn that is more prone to aggregation. H4 cells were transiently transfected with constructs encoding for Synphilin-1 and SynT, as previously described (Lázaro et al, 2014), in order to induce the formation of LB-like inclusions (Supplementary figure S2A), as characterized by our and other groups (Engelender et al, 1999; Masaracchia et al, 2020; McLean et al, 2001). Seventeen hours after transfection, cells were treated with doxycycline at the following concentrations: 0.01 μM, 0.1 μM, 1 μM and 10 μM. The presence and number of inclusions were assessed by immunocytochemistry (Figure 3A) 24 hours after treatment. An average of 50 positive cells for aSyn expression per treatment and experiment were then arbitrary divided into four different groups, depending on the number of inclusions present, i.e. cells without inclusions, cells with 1 to 5 inclusions, cells with 6 to 10 inclusions, and cells with more than 10 inclusions (Figure 3 A-E). We observed no statistically significant differences in the percentages of transfected cells showing less than six inclusions with any of the doxycycline concentrations tested (Figure 3 C). On the other hand, the percentage of cells containing more than 6 inclusions significantly decreased when cells were treated with 10 μM doxycycline (Figure 3 D, E). Interestingly, the percentage of cells showing more than 10 inclusions increased when cells were treated with 0.01, 0.1 and 1 μM. The total number of inclusions per cell showed a tendency to increase in relation with the control cells at 0.1 and 1 μM, with values of 4.05 ± 1.57, 3.97 ± 1.41, 5.86 ± 0.88 and 6.08 ± 1.56 inclusions per cell, respectively (Supplementary figure S2B). Treatment with 10 μM resulted in a reduction in the total number of inclusions per cell (1.28 ± 0.39 inclusions per cell).

**Figure 3.**
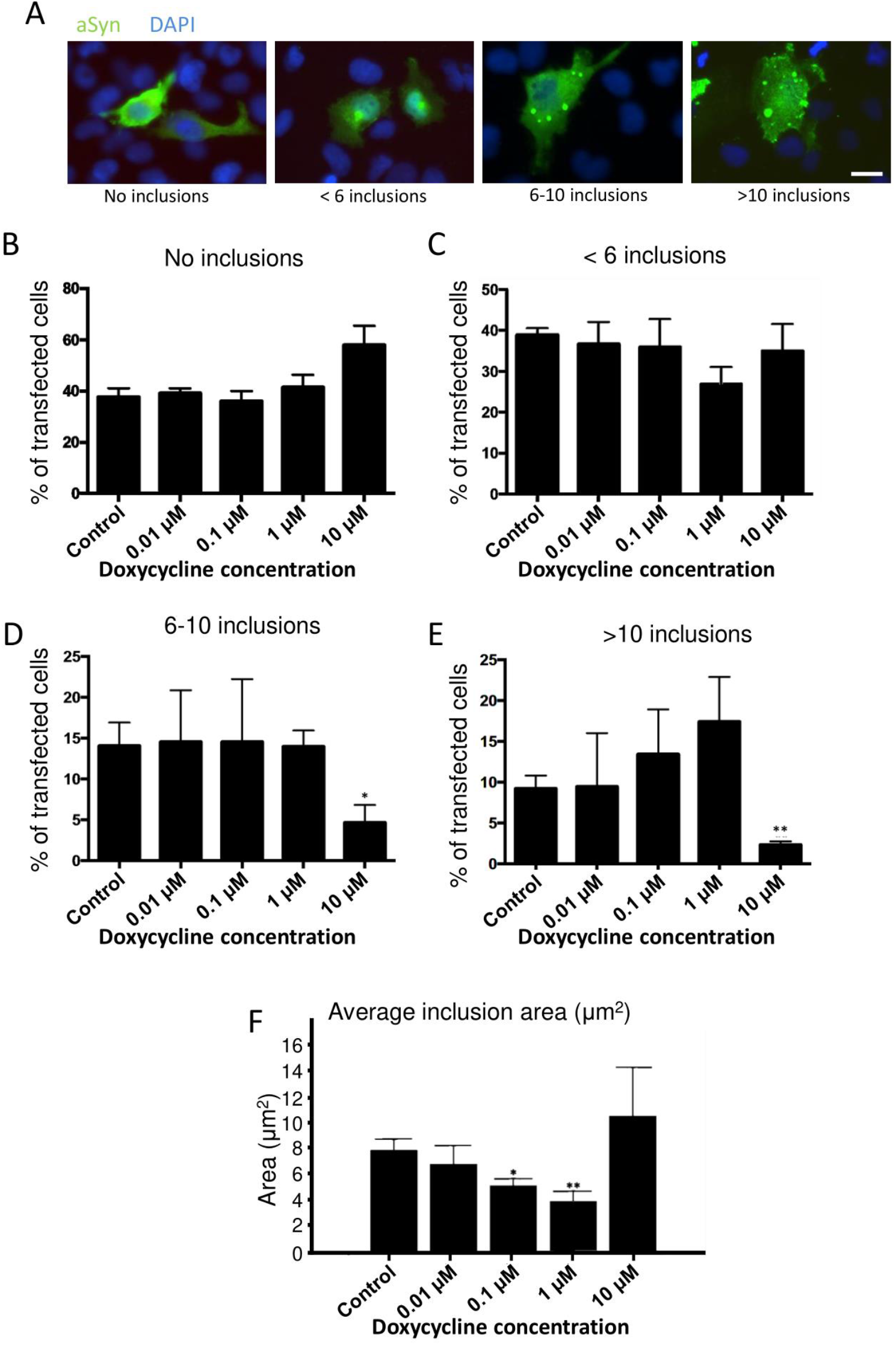
Doxycycline reduces aSyn inclusion size and number in H4 cells. **A.** Representative pictures of immunostained H4 cells without inclusions, with less than six inclusions, between six and ten inclusions and with more than 10 inclusions. Scale bar 25 μm. **B.** Percentage of cells that showed no inclusions in the absence or in the presence of doxycycline (0.01-10 μM). **C**. Percentage of cells that showed less than six inclusions in the presence or not of doxycycline (0.01-10 μM). **D.** Percentage of cells that showed between six and ten inclusions in the absence or presence of doxycycline (0.01-10 μM). Treatment with 10 μM doxycycline leads to a significant decrease in the percentage of transfected cells containing between 6 and 10 inclusions. **E.** Percentage of cells that showed more than ten inclusions over the total number of transfected cells in the absence or presence of doxycycline (0.01-10 μM). There was a significant decrease in the percentage of cells treated with 10 μM doxycycline. **F.** Average area of the inclusions in cells positive for aSyn expression at different doxycycline concentrations. Treatment with 0,1 and 1 μM doxycycline leads to a significant decrease in the percentage of transfected cells that showed more than ten inclusions. Data presented as mean ± SD. N=3 in all experiments * p < 0.05 relative to control. ** p < 0.005 relative to control. ANOVA and subsequent t-test against control.

### Doxycyline reduces the size of aSyn inclusions

After observing a slight increase in the percentage of cells showing more than 10 inclusions (Figure 3E), we wondered whether this was due to the formation of smaller inclusions. To assess this, we checked the average inclusions size. Untreated cells showed average values of 7.85 ± 0.82 μm^2^ and cells treated with 0.01 μM of doxycycline showed average values of 6.80 ± 1.42 μm^2^. Interestingly, we observed a significant decrease in average inclusion area when cells were treated with 0.1 and 1 μM, values been 5.13 ± 0.56 and 3.91 ± 0.79 μm^2^, respectively (*p* = 0.03 for 0.1 μM, n = 3 and *p* = 0.004 for 1 μM, n = 3). This was reverted when cells were treated with 10 μM doxycycline, as the mean area was 10.50 ± 3.72 μm^2^ (Figure 3F). The greater average size of inclusions in cells treated with 10 μM combined with the lower average number of inclusions may suggest that, at 10 μM doxycycline, smaller inclusions are disrupted while a few larger ones still remain. Furthermore, the total inclusion area in the cells was also smaller, which points to not only a disruption of smaller inclusions but to an overall lower presence of aggregated aSyn (Supplementary figure S2C).

To test whether doxycycline was disrupting the aggregates into smaller ones, we performed live cell imaging of cells transfected with aSyn-VC and VN-Synphilin-1 bimolecular fluorescence complementation constructs (Lázaro et al, 2014) and manuscript in preparation). As the inclusions were formed through the interaction between aSyn and Synphilin-1, the VN- and VC-fragments of Venus fluorescent protein came into contact, thereby reconstituting the functional fluorophore and emitting fluorescence (Outeiro et al, 2008). Using this aggregation model we observed a decrease in particle size over time after treatment with 10 μM doxycycline (Supplementary figure S2D), consistent with the disruption of aSyn inclusions.

### Doxycycline prevents the seeding of aSyn inclusions in PFF-treated cells

It has been reported that the addition of exogenous aSyn PFFs seeds the aggregation of endogenous aSyn (Volpicelli-Daley et al, 2014). Thus, based on our previous results, we hypothesized that doxycycline could inhibit this process. To assess this we treated, 48 hours after plating, cells stably expressing aSyn-EGFP with PFFs at a final concentration of 100 nM of PFFs with 1 or 10 μM doxycycline. We observed a significant decrease in the number of inclusions in cells treated with 10 μM doxycycline (Figure 4A, B) when compared to vehicle-treated cells.

**Figure 4.**
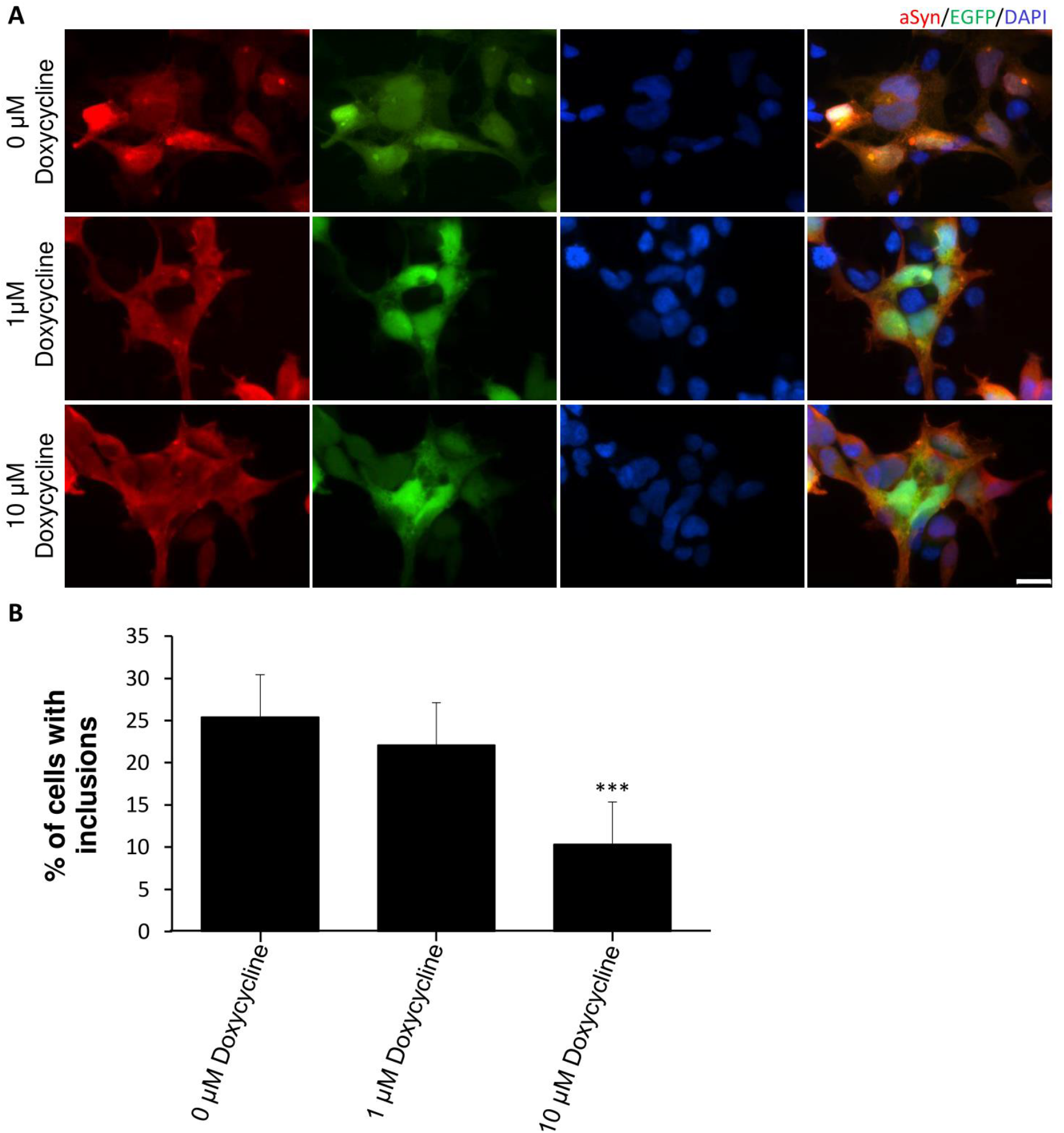
Doxycycline prevents the formation of aSyn inclusions after treatment with exogenous PFFs. Treatment with PFFs at a concentration of 0.1 μM for 48hrs in absence or presence of doxycycline at concentrations of 1 μM and 10 μM. **A** Presence of aSyn aggregates in HEK293-aSyn:EGFP stable cells treated with PFF and visualized with anti-aSyn antibody (Scale bar 20 μm). DAPI was used for nuclear staining. **B**. Quantification of the inclusions formed in n>200 cells per condition (mean ± SD) reveals a significant decrease in the number of inclusions after treatment with 10 μM doxycycline. In contrast, treatment with 1 μM of doxycycline had no effect compared to non-treated cells. N = 3 independent experiments ** p < 0.005.

### Expression of human aSyn in the dopaminergic neurons of *C. elegans*

We first characterised the presence of the aSyn in the tagged dopaminergic neurons. For this, we evaluated the specific targeting pattern of the *dat-1p* promoter using GFP as a reporter. We observed signals in all the eight DA neurons previously described (Sulston et al, 1975), including (i) the four CEP cell bodies, their associated neuritic extensions and the two ADEs neurons located in the anterior part of the animals, and (ii) the two PDEs neurons in a more posterior lateral position (Figure 5A). Using an anti-pan aSyn antibody, we confirmed that the aSyn transgenic line expressed aSyn in all GFP^+^ DA neurons (Figure 5B). In addition, GFP^+^ neurons were also immuno-positive for TH. (Fig 5B)

**Figure 5.**
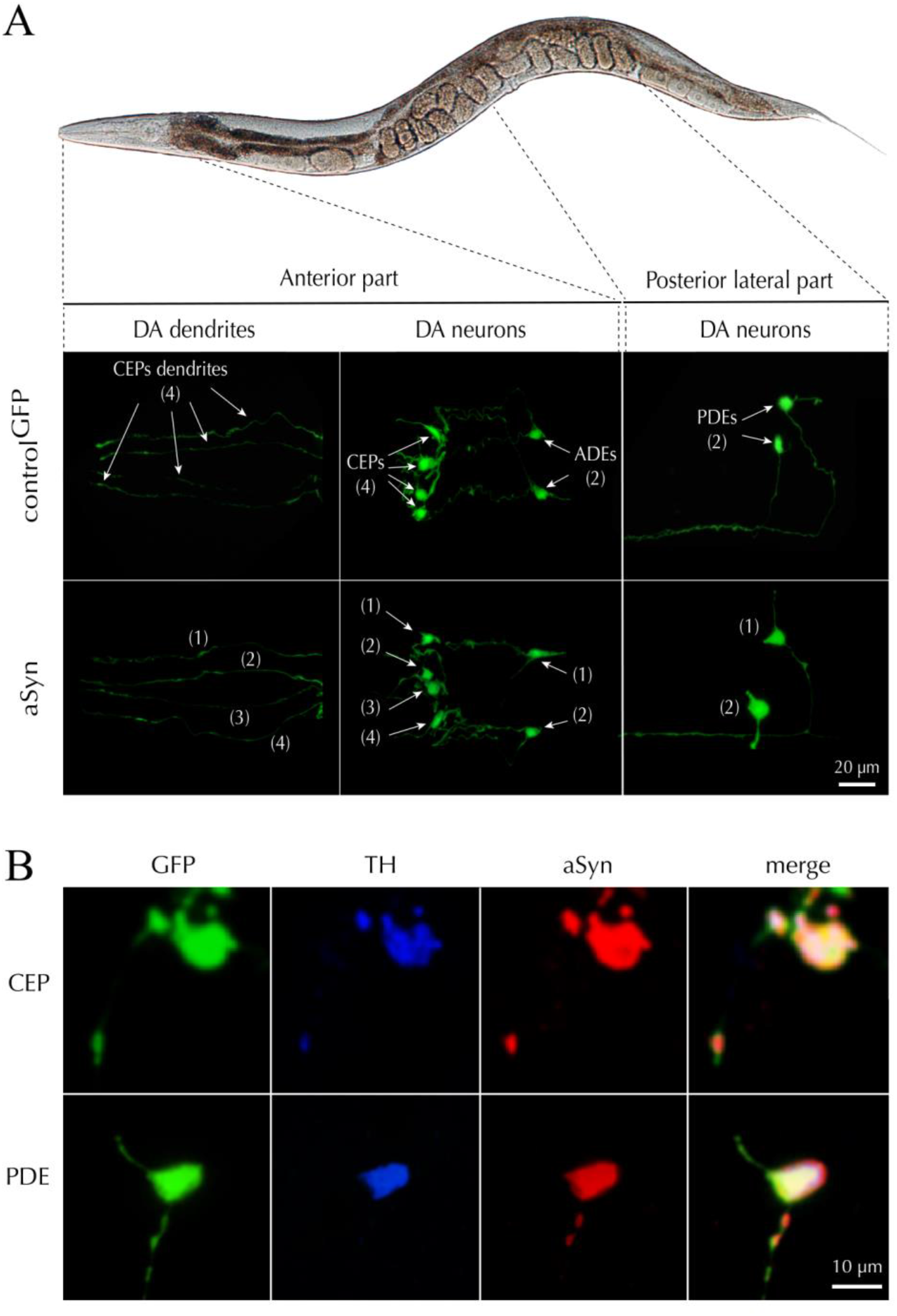
Targeted expression of aSyn in *C. elegans* DA neurons. **A.** Illustration describing a *C. elegans* control line expressing the GFP reporter gene (control^GPF^) specifically in DA neurons and another line co-expressing the wild-type human *SNCA* gene and the GFP reporter (αS), in the same neurons. The observation was made on fixed samples using an epifluorescence microscope fitted with a high magnification (x100) objective. The micrographs depict the expression pattern of the GFP reporter in the two lines and confirm the presence of six DA neurons in the head (CEPs and ADEs) and two DA neurons (PDEs) in the middle of the body. **B.** Panel showing the concomitant expression of TH (blue), aSyn (red) and GFP (green) in the CEP and PDE neurons. These observations confirm that aSyn is expressed in the soma and the dendrites of DA neurons.

### Doxycycline alters the subcellular distribution of aSyn in dopaminergic neurons in *C. elegans*

The total number of DA neurons tagged with GFP was similar in animals expressing aSyn and controls suggesting that the expression of aSyn does not induce DA neuronal loss. However, the expression of aSyn induced both a significant increase in the surface area of the DA cell bodies of 60.58% (54.54 ± 3.44 μm^2^ for control^GFP^ versus 87.58 ± 7.73 μm^2^ for aSyn line; *p* = 0.0004) and a significant decrease of GFP density of 23.87% (118.5 ± 12.80 A.U. for control^GFP^ versus 90.21 ± 5.04 A.U. for aSyn line; *p* = 0.03) suggesting a deleterious effect of aSyn expression on DA neurons (Figure 6A). Chronic treatment with 15 μM doxycycline did not induce any change in the number of DA cell bodies or their neuritic processes in comparison to untreated controls. In contrast, this tetracycline concentration significantly decreased the swollen morphology of the soma induced by the aSyn transgene. In addition, doxycycline significantly improved the altered density of GFP fluorescent signal induced by aSyn when comparing to untreated animals (Figure 6A). In addition, we performed immunohistochemistry of aSyn on whole animal observing a regular immunolabeling of the aSyn in the soma and along the axons of the PDEs GFP tagged neurons (Figure 6B, left panel). The treatment with doxycycline altered the distribution of the protein in the soma compartment, presenting a punctuated expression pattern (white arrows) (Figure 6B, left panel). To evaluate these modifications, we quantified the ratio of the surface area signal between the aSyn associated immunofluorescence and the GFP signal in the soma of PDE neurons. In the presence of doxycycline, we found a significant reduction of 55.20% f the surface area of the aSyn distribution signal relative to the soma of the PDE neurons (Figure 6B, right panel).

**Figure 6.**
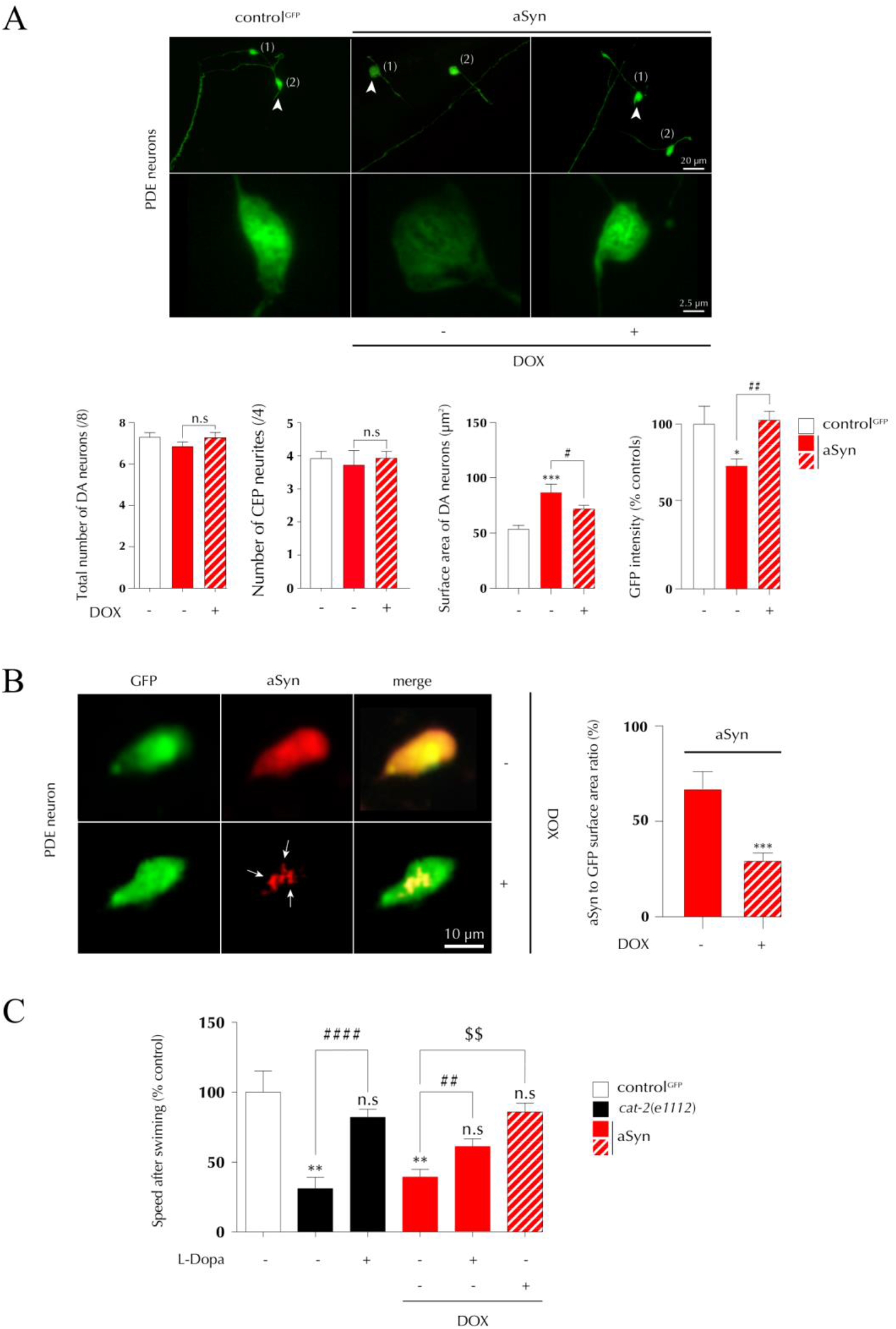
Cellular and behavioural consequences of doxycycline treatment in *C. elegans*. **A.** After synchronization, a homogeneous population of young adult animals treated or not with doxycycline (15 μM), were PFA-fixed (4%), mounted on slide and observed with an epifluorescence microscope at equipped with a x63 objective. (Upper panel) Images illustrate the most characteristic features of DA GFP-tagged neurons (PDE, arrows head). (Lower panel) Quantification of the number of GFP^+^ cell bodies and dendrites, the surface area of GFP+ cell bodies and the somal intensity of the GFP fluorescent signal in control and aSyn-expressing animals. Bar graphs show that aSyn expression did not affect the number of DA neurons but induced significant morphological alterations, such an increase in the somal surface area and a decrease of the GFP fluorescence signal. Such changes were prevented by a chronic exposure to doxycycline. Data are expressed as mean values ± s.e.m. Three independent experiment; *n* = 25/condition/experiment. Statistical significance was calculated with various tests: (i) a oneway ANOVA followed by a *Dunnett’s* multiple comparisons tests with all condition versus the untreated control line (control^GFP^), \**p*<0,05 and \*\*\**p*<0,005; (ii) a two-tailed unpaired *t-test* for aSyn animals treated or not with doxycycline, ^#^*p*<0,05, and ^##^*p*<0,01; n.s: not significant. **B.** *Left panel*: Illustrations showing the impact of doxycycline (15 μM) on the distribution of human aSyn in the soma of DA neurons. Representative images of the wild-type human aSyn immunosignal (red signal) in the PDE-DA GFP tagged neurons (green). In untreated animals the immunosignal for aSyn is distributed over a large part of the cell body in contrast with worms treated with doxycycline where the signal is punctuated and restricted to a small intracellular compartment (white arrows). *Right panel*: Measurement of the impact of the doxycycline treatment on the surface area occupied by the aSyn immunosignal in the soma of PDE DA neurons. The surface area of the aSyn immunosignal expressed as a ratio to the total somal surface area of GFP positive neurons, appears significantly decreased. Data are mean values ± s.e.m. Three independent experiment, *n* = 20/condition/experiment. Statistical significance was calculated with two-tailed unpaired *t-test* for the two experimental conditions; \*\*\**p*<0,005. **C.** Impact of the doxycycline treatment on behavioural crawling velocity after swim-to-crawl transition. Note that the loss of velocity in animals expressing wild-type aSyn is significantly prevented by a chronical doxycycline treatment in comparison with the untreated line. The null mutant *cat-2 (e1112)* /TH was used as positive control for the assays. A treatment of the animals with 1 mM of L-DOPA 2h before the test prevented the behavioural defect observed in the line expressing aSyn. These results suggest that DA neurons are affected by the presence of the aSyn. Data are mean values ± s.e.m. Three independent experiment, *n* = 30/condition/experiment. Statistical significance was calculated with various tests: (i) a one-way ANOVA followed by a *Dunnett’s* multiple comparisons test versus the GFP control line (Control^GFP^); \*\**p*<0,01, n.s is not significant; (ii) a two-tailed unpaired *t-test* for *cat-2 (e1112)* animals treated or not with L-Dopa, ^####^*p*<0,001; (iii) a two-tailed unpaired *t-test* for aSyn animals treated or not with L-Dopa, ^##^*p*<0,01; (iv) a two-tailed unpaired *t-test* for aSyn animals treated or not with doxycycline, ^$$^*p*<0,01.

### Doxycycline restores motor alteration in aSyn *C. elegans*

The functionality of DA neurons was evaluated using a specific dopaminergic behavioural test which measure the locomotor velocity of the animal in a plate preceded with a short swimming activity in a liquid media (Vidal-Gadea et al, 2011). As expected, we found that this parameter was significant decreased in the *cat-2* (*e1112*) TH null mutant used as a reference line (Figure 6C). Most interesting, this parameter was strongly reduced in the transgenic line expressing aSyn in DA neurons in comparison to the control line (Figure 6C). L-Dopa treatment significantly restored dopaminergic function in both the *cat-2* and aSyn lines. These results suggest that aSyn overexpression in the DA neurons resulted in a DA deficit inducing behavioural dysfunction despite the absence of neuronal loss. Of interest, doxycycline treatment restored locomotor behaviour specifically on aSyn line (Figure 6C) compared to the control strain expressing only GFP (not shown), suggesting that the effect of doxycycline on aSyn distribution in DA neurons improved motor behaviour.

## Discussion

The aggregation of aSyn is one of the hallmarks of synucleinopathies. In recent years, several aggregation inhibitors have been developed and tested, and the search for new ones continues (Dominguez-Meijide et al, 2020; Hengartner & Fernandez, 2019). Different drugs have been developed and tested, such as pyrazole derivatives and hydrazones (Cukierman et al, 2020; Cukierman et al, 2017; Wagner et al, 2013). Nonetheless, most of the drugs developed are either toxic or show low bioavailability in the brain due to low water solubility or other factors. To circumvent this, new therapeutic strategies like pharmacokinetic modulation (like complexation with cyclodextrines) or repurposing of existing drugs have been developed (Barros et al, 2018; Heemskerk et al, 2002). Examples of recent repurposing of existing drugs include inhibitors of the renin-angiotensin system, anti-asthmatic drugs, COX-2 inhibitors and other NSAIDs (Hengartner & Fernandez, 2019; Jang et al, 2017; Moghaddam et al, 2007; Rodriguez-Perez et al, 2018). Following this trend, the anti-neuroinflammatory properties of tetracyclines such as minocycline and doxycycline have been studied (Bortolanza et al, 2018; Lazzarini et al, 2013). More recently, it has been proposed that tetracyclines and, specifically, doxycycline, have not only anti-neuroinflammatory properties, but can also inhibit the aggregation of misfolded aSyn (González-Lizárraga et al, 2017).

Our results in cell free systems showed that increasing concentrations of doxycycline lead to a decrease in ThT fluorescence intensity in an RT-QuiC assay. This is probably due not only to a decrease in the aggregation rate of aSyn, but also to a possible change in the conformation of the aSyn aggregates formed. As ThT becomes fluorescent upon binding to beta-sheet structures, a possible change in aSyn aggregation may lead to lower ThT binding and, thereby, to lower ThT fluorescence (Wu et al, 2009). Indeed, it has been proposed that treatment with doxycycline induces a conformational change, as infrared studies demonstrated the presence of parallel beta-sheet instead of the typical antiparallel beta-sheet found in on-pathway toxic oligomeric species (Celej et al, 2012). Furthermore, our WB, EM and DLS results confirm previous observations where it was reported that the presence of doxycycline induces a remodelling of aSyn oligomers into off-pathway non-toxic, non-seeding, high molecular weight species (González-Lizárraga et al, 2017). Our results and previous ones also prove that this effect is dose-dependent (González-Lizárraga et al, 2017). In addition, our RT-QuiC results show a decrease in ThT fluorescence intensity at high doxycycline concentrations. As these reactions were seeded, this suggests additional effects on the fibrils. Furthermore, we also demonstrate that while aSyn aggregates are very efficient at seeding native aSyn, as revealed by ThT assays, they lose all seeding ability when incubated with doxycycline.

Our results in cells also show the ability of doxycycline to break intracellular aSyn-positive inclusions, as we observed an increase in the number inclusions per cell that was accompanied by a decrease in their size. This was further confirmed by live cell imaging. The lower percentage of cells with more than ten inclusions when cells were treated with 10 μM, and the higher average inclusion size may be either due to more compact inclusions that cannot be decomposed by doxycycline or to a possible reorganization of aSyn inside individual cells that leads to fewer but larger inclusions. In addition, it has been proposed that exogenous aSyn fibrils can be internalized and act as templates for the formation of aggregates by addition of endogenous monomeric aSyn (Grassi et al, 2018; Masaracchia et al, 2018; Volpicelli-Daley et al, 2014). These results, in combination with those observed in cell free systems, further support the idea of doxycycline being able to exert inhibition of seeding. Thus, doxycycline not only degrades already formed aggregates, but it can also inhibit the formation of new ones. It has been reported that an equimolar concentration of doxycycline to aSyn would be suitable to reach a protective effect (González-Lizárraga et al, 2017). The concentration of 10 μM may be already above equimolar levels as the concentration of aSyn in cell synapse is ~50 μM (Perni et al, 2017; Wilhelm et al, 2014). Furthermore, the neuroprotective and anti-inflammatory doses of doxycycline are below the ones needed to exert its antibiotic activity (González-Lizárraga et al, 2017; Santa-Cecília et al, 2019).

An additional pathological effect of aSyn does not come from its aggregation and propagation. It may also cause mitochondrial dysfunction and enhanced production of ROS, which may, in turn, further modulate aSyn oligomerization and cytotoxicity, leading to a vicious circle. The generation of ROS by aSyn aggregates can affect macromolecules or activate signalling pathways that culminate in central nervous system neurodegeneration (Park et al, 2020; Wang et al, 2019). For this reason, the inhibition of oxidative stress could offer an effective therapeutic approach to mitigate the progression of PD. In concordance with this concept, we show that aSyn aggregates increase the production of ROS in differentiated SH-SY5Y cells. However, aSyn aggregates formed in presence of doxycycline are less prone to induce oxidative stress in cell cultures. In this regard, minocycline and doxycycline have been proposed to protect against mitochondrial dysfunction. Furthermore, the proposed mechanism of inhibition of aSyn aggregation may also reduce its detrimental effects on mitochondrial function (Garcia-Martinez et al, 2010; Santa-Cecília et al, 2016).

In recent years, several aSyn-inducible cell models have been developed, including tet-on and tet-off inducible systems, which are based on the induction or blocking of aSyn expression in the presence of doxycycline, respectively (Vekrellis et al, 2009; Zhu et al, 2002). These induction systems have been developed in several cell lines and animal models (Harding et al, 1997; Takahashi et al, 2007; Vasquez et al, 2018; Vekrellis et al, 2009). In tet-on systems, addition of doxycycline leads to the expression of aSyn at levels that enable the formation of inclusions (Vasquez et al, 2018). The concentrations of doxycycline used in these publications range between 2.25 and 6.7 μM (Harding et al, 1997; Vasquez et al, 2018). Our results show a stronger effect above equimolar concentrations. Therefore, the concentration of doxycycline required to modulate protein aggregation is higher than that required for inducing aSyn expression.

To further test the biochemical properties of doxycycline in this context, we generated a new *in vivo* transgenic model of PD in *C. elegans* expressing wild-type aSyn in the DA neurons. Unlike previously established *C. elegans* models designed to model different features of PD (Cooper & Van Raamsdonk, 2018), and target tissues like muscle (Hamamichi et al, 2008; van Ham et al, 2008), the entire neuronal system (Kuwahara et al, 2006; Kuwahara et al, 2008; Lakso et al, 2003), or the DA system (Cao et al, 2005; Lakso et al, 2003), our model uses the *dat-1* promoter. Dat-1 is the ortholog of the human dopamine transporter in the nematode, and is selectively expressed in the DA system. The model enables the *in vivo* study of the specific vulnerability of the DA neurons due to the presence of aSyn. We verified the correct targeting of the transgene in the cell soma using GFP as a reporter. Human wild-type aSyn, co-expressed with the GFP reporter under the same genetic regulatory system, is present in the soma of the DA neurons. In young-adult animals, aSyn did not induce significant DA neuronal loss until the worms reached 6 days of age. In another *C. elegans* model expressing aSyn in the DA system, the authors described the presence of aSyn-dependent neuronal injuries, probably resulting from differences in the expression levels of the transgene (Cao et al, 2005). However, in our model aSyn induced abnormal neuronal morphologies consistent with a dysfunctional DA system, as indicated by a noticeable alteration of DA-associated behaviour, suggesting that overexpression of aSyn impacts DA homeostasis. These irregular neuronal morphologies and pathological behaviour are restored after chronic treatment with doxycycline, consistent with the effects observed *in vitro* model, and reinforcing the hypothesis that doxycycline may act as a neuroprotective agent.

It has been proposed that *C. elegans* movement depends not only on DA, but also on cholinergic neuron-dependent muscle contraction and GABAergic neuron-dependent relaxation. However, it has been shown that blockade of DA D2 receptors does not fully limit paralysis triggered by changes in cholinergic and GABAergic signaling (Refai & Blakely, 2019). The opposite results have also been shown, i.e. changes in cholinergic and GABAergic signaling due to D2 antagonism (Krum et al, 2020). Furthermore, it was also shown that changes in DA may affect GABAergic egg release in the worms (Cermak et al, 2020). These results demonstrate that further research into these pathways is necessary, but this is beyond the scope of our study.

In our study, we employed a classic transgenesis technique (Mello & Fire, 1995) to produce our line, resulting in the production of transgenic animals carrying multi-copy arrays of the *SNCA* transgene, which may induce non-specific neurobiological process. Additionally, some studies showed that expression of aSyn in DA motor neurons *of C. elegans* recapitulates features of aSyn neuronal propagation. Using the BiFC assay (Outeiro et al, 2008), we showed the protein could disseminate to the neighbouring serotoninergic neurons and interact with another host targeted aSyn (Tyson et al, 2017). These processes have been extensively described in mammals for synucleinopathies and other proteinopathies (Peng et al, 2020). Here, we did not observe a significant neuronal propagation of aSyn, which might be possibly explained by the fact that we used a different promoter gene to induce aSyn expression. In addition, we have not investigated the tissue expression pattern of aSyn in old animals, as this was beyond the scope of the present study.

In *C. elegans* the treatment with doxycycline induces a significant effect on development by reducing the growth rate (Letizia et al, 2018), and it appears to promote longevity via inhibition of *mrps-5* ribosomal protein and atf-4 transcription factor mitochondrial pathways (Molenaars et al, 2020). Doxycycline has been described to play role in the inhibition of the translation process effect by suppressing the binding of tRNAs to the ribosomal-machinery system and inducing a global decrease in protein production (Nelson & Levy, 2011). Thus, we cannot rule out that the neuroprotective effect of doxycycline in our experimental conditions may also be explained *via* an indirect activation of the aging-metabolic cellular pathways and the observed effect on mitochondria and ROS levels, as decreased ROS may lead to a lower level of neuroinflammation and neuroprotective effects (Picca et al, 2020). Since aging is the major risk factor for PD, it will be interesting to assess the potential interaction between aging-associated pathways and aSyn in the novel transgenic lines we generated.

Strikingly, we observed modifications in aSyn distribution in the DA neurons in aSyn-transgenic animals chronically incubated with doxycycline. In particular, we observed a reduction of the sub-cellular distribution of the protein in the soma. This occurred together with the effects on aSyn aggregation, further strengthening the anti-aggregation properties of doxycycline *in vivo*, and suggesting additional complementary effects on aSyn to those observed *in vitro* and in cell-free models. The punctate pattern observed in DA neurons *in vivo* are consistent with a combined effect of changes in cellular distribution and the formation of fewer and larger inclusions inside the cells, as observed *in vitro*. These changes lead to an overall decrease in aSyn area in relation to the total surface of the cell soma of the PDE neurons. This is not an independent effect, but the combination of different effects, i.e. inhibition of aggregation, disruption of pre-formed inclusions, blocking of seeding, and changes in inclusion area leading, ultimately, to the distinct dotted pattern observed *in vivo*. Additionally, aggregated species of aSyn produced in the presence of doxycycline lost their capacity to stimulate mitochondrial ROS production, which may also account for the restoration of DA neuron function. In fact, our results indicate that a restoration of motor behavior by doxycycline can be explained by the combination of the aforementioned decrease in ROS, combined with changes in aSyn processing, seeding and distribution.

As mentioned above, doxycycline has neuroprotective effects in several models of neurodegenerative disease and, in particular, has anti-aggregation properties (Costa et al, 2011; Paldino et al, 2020; Santa-Cecília et al, 2019). In a *C. elegans* amyloidogenesis model induced by the expression of a human pathogenic variant of β2-microglobulin (D76N of β_2_-m) in the muscle (Faravelli et al, 2019), doxycycline decreases the proportion of oligomeric fraction produced by the β_2_-m aggregation, resulting in an improvement of the motor symptoms (Raimondi et al, 2017). These findings are consistent with those we report herein.

In summary, our data provides novel insight into the mechanisms by which doxycycline may inhibit the aggregation of aSyn by remodelling aSyn oligomers into off-pathway non-toxic, non-seeding, high molecular weight species. We showed this effect not only *in vitro* but also in cell free models. The extreme need for alternative strategies against PD calls for attempts for repurposing existing drugs. In this context, our data suggests that doxycycline may act at different levels, modulating the aggregation process and the effects of aSyn aggregation, promoting somatic repair, and leading to motor recovery in *C. elegans*.

## Materials and methods

### Cell-free studies

#### Generation of pre-formed fibrils

Monomeric aSyn WT was produced by transforming *E. coli* BL21-DE3 competent cells with plasmids encoding the corresponding cDNA sequence (pET21-aSyn). Purification was performed as previously reported (Masaracchia et al, 2018; Volpicelli-Daley et al, 2014) and the protein stocks were frozen in single aliquots at −80°C. The stocks were subsequently diluted in a buffer containing 100 mM Tris, 500 mM NaCl and 10 mM EDTA to a final concentration of 5 mg/mL. Samples were incubated for seven days in an Eppendorf Thermomixer Comfort (Eppendorf, USA) at 600 rpm and 37°C. The transition from the monomeric form to aggregated aSyn was determined by thioflavin T fluorescence absorbance. In brief, thioflavin T was added to a final concentration of 20 μM, samples were mixed by slowly inverting the tubes and the mix was added to a black 96-well plate (COSTAR, Corning Incorporated) in which 100 μl of the reaction mixture was loaded. Fluorescence intensity measurements were performed in an Infinite M200 fluorescence plate reader (TECAN) at 480 nm. Three replicates of each sample were measured.

#### Seeding assay

Monomeric aSyn solutions (70 μM) in 20 mM HEPES, 150 mM NaCl, pH 7.4, were incubated in a Thermomixer comfort (Eppendorf) at 37°C under orbital agitation at 600 rev./min to form aggregates. These species were diluted 1/10 when harvested and further incubated with 70 μM of fresh monomers in the absence or presence of 100 μM doxycycline. Aliquots were taken from the seeding aggregation at different times and mixed with ThT probe, according to LeVine (LeVine, 1993). Changes in the emission fluorescence spectra with the excitation wavelength set at 450 nm were monitored using FluoroMax-4 Spectrofluorometer. We employed similar proportions between oligomers, monomers and ThT as the ones used for the RT-QuiC.

#### Real-time quaking induced conversion (RT-QuiC)

RT-QuiC reactions were performed in black 96-well plates (COSTAR, Corning Incorporated) in which 100 μl of the reaction mixtures were loaded. The final concentrations in the reaction mix were 150 mM NaCl, 1 mM EDTA, 10 μM ThT, 70 μM SDS, 7 μM monomeric aSyn and 0.7 μM of PFF in PBS buffer (pH= 7.1). Doxycycline was added at the indicated final concentrations. Plates were covered with sealing tape and incubated at 41°C in a plate reader (Infinite M200 fluorescence plate reader, TECAN) with intermittent shaking cycles, consisting of one minute orbital shaking at 432 rpm followed by two minutes incubation and one minute pause to measure the ThT fluorescence intensity at 480 nm. Three replicates of each sample were measured for 250 amplification cycles. Pre-formed fibrils, generated by the method proposed by Volpicelli-Daley have been shown to have similar aggregation and spreading behavior to in vivo generated seeds (Patterson et al, 2019; Volpicelli-Daley et al, 2014), furthermore, as in vivo generated seeds need to be purified from patients there are several external factors affecting the homogeneity of the results including the baseline pathology (Candelise et al, 2019; Candelise et al, 2020). Thus, the use of in vitro generated seeds will lead to more comparable results.

#### Dynamic light scattering

Dynamic light scattering measurements were performed using a DynaPro Nanostar (Whyatt) operating at λ = 633 nm. The measurement temperature was 25°C. Protein samples contained 7.7 μM of aSyn in RT-QuiC aggregation buffer. Measurements were always performed in triplicate.

### *In vitro* studies

#### Cell culture

Human neuroglioma (H4) cells were maintained in OptiMEM 1x with Glutamax medium (Gibco, Carlsbad, Germany) supplemented with 10% FBS and 1% penicillin/streptomycin (PS) at 37°C in a 5% CO_2_ atmosphere.

Human embryonic kidney (HEK293) cells were maintained in DMEM supplemented with 10% FBS and 1% PS at 37°C in a 5% CO_2_ atmosphere.

Human neuroblastoma (SH-SY5Y) cells were used to measure reactive oxygen species produced by mitochondria. Cells were grown in DMEM-GlutaMAX supplemented with 10% FBS and 1% PS, at 37 °C and 5% CO2. After, the cells were seeded in 48 plates at 60,000 cell/well and differentiated 24 h after plating by treatment with 10 μM retinoic acid (Sigma) in DMEM-GlutaMAX supplemented with 1% FBS and 1% PS, at 37 °C and 5% CO2 for 5 days. Media were changed every 2 days.

#### Cell transfection

Twenty-four hours prior to transfection, approximately 100.000 H4 cells were plated per well in a 12-well plate (Costar, Corning, New York, USA). Six hours prior to transfection medium was replaced with fresh one. Five μl of a 2.5 M CaCl_2_ solution was added to 45 μl of a solution containing equal amounts of the plasmids encoding SynT and Synphilin-1-V5 (Lázaro et al, 2014). 2xHBS calcium phosphate buffer (50 mM N, N-bis (2-hydroxyethyl)-2-aminoethanesulfonic acid (BES) pH 7.05, 280 mM NaCl, 1.5 mM Na_2_PO_4_ x 2 H_2_O), pH 7.05 was added drop wise to the solution and vigorously mixed. The mix was incubated for 20 minutes and added dropwise to the cells while the plate was gently rocked.

Additionally, HEK293 cells were transfected according to the following protocol: twenty-four hours prior to transfection, approximately 50.000 cells were plated per well in a 24-well live imaging plate (Zell-Kontakt, Nörtern-Hardenberg, Germany). Transfection was performed with metafectene according to the manufacturer’s instructions. In brief, a solution containing equal amounts of the plasmids encoding aSyn-VC and VN-Synphilin-1 (Lázaro et al, 2014) was added to a solution containing metafectene leading to a final DNA:metafectene ratio of 1:3.

#### Cell treatment

Seventeen hours after transfection, H4 cells were fed with fresh medium and treated with doxycycline hyclate (Sigma-Aldrich, St. Louis, MO, USA) at the indicated final concentrations. Additionally, some cells were treated with water as it was used as solvent for doxycycline.

Seventeen hours after transfection, HEK293 cells were fed with fresh medium and treated with doxycycline hyclate to a final concentration of 10 μM.

Additionally, exogenous pre-formed fibrils (PFFs) (Volpicelli-Daley et al, 2014) were added to HEK293 cells stable for the expression of aSyn-EGFP at 0.1 μM 48 hours after culture. Concomitantly, 1 and 10 μM doxycycline were added to the cells. At the end of the treatment, cells were extensively washed with PBS to remove residual proteins still outside of the cells and immunocytochemistry was performed.

#### Mitochondrial superoxide levels

Differentiated SH-SY5Y cells were treated with aggregates produced in the absence or presence of 100 μM doxycycline. ROS levels produced by cells were examined by determining mitochondrial superoxide using MitoSOX Red. This probe selectively targets mitochondria and undergoes oxidation by superoxide to form a stable fluorescent compound. Briefly, SH-SY5Y cells after the different treatments were incubated with 5 μM MitoSOX Red reagent (Invitrogen) for 10 min, then washed and fixed with 4% formaldehyde in PBS before further analysis. Fluorescent images of randomly chosen fields were acquired with identical acquisition parameters using inverted fluorescence microscopy. The experiments were run in triplicate and fluorescence intensity values calculated by adding the intensities of each pixel in the image and then the average background was subtracted.

#### Immunoblotting

For SDS-PAGE, a volume of the RT-QuiC products containing 0.2 μg of aSyn was boiled with Laemmli buffer at 95°C for 5 minutes and loaded into 12% Bis-Tris-polyacrylamide gels and transferred to nitrocellulose membranes. The membranes were incubated overnight with different anti-aSyn primary antibodies with different binding properties: either 1:1000 mouse anti-aSyn (BD Biosciences), 1:3000 mouse anti-aggregated-aSyn (clone 5G4, Millipore), 1:2000 rabbit anti C20-Synuclein (Santa Cruz) or 1:2000 rabbit anti-αβγSyn (Santa Cruz). For the SDD-PAGE immunoblot, the method used was a modification of a previously proposed one (Olsson et al, 2018). In brief: a volume of the RT-QuiC products containing 0.2 μg of aSyn was loaded into 10% tricine gels. The electrophoresis was performed with a running buffer containing 0.1 M Tris base, 0.1 M tricine and 0.1% SDS. Gels were then transferred to nitrocellulose membranes. The membranes were incubated overnight with 1:3000 mouse anti-aggregated-aSyn (clone 5G4, Millipore) at 4°C. Following three washes with TBS-Tween, membranes were incubated for two hours with the corresponding HRP conjugated secondary antibody (GE Healthcare). After incubation with the secondary antibody, membranes were washed three times with TBS-Tween and developed in a chemiluminescence system (Fusion FX Vilber Loumat).

#### Immunocytochemistry

Twenty-four hours after treatment with doxycycline, transfected H4 cells were washed with PBS and fixed with 4% PFA for 20 minutes at room temperature (RT), followed by a permeabilization step with 0.1% Triton X-100 (Sigma-Aldrich, St. Louis, MO, USA) for 20 minutes at RT. Following permeabilization, a blocking step with 1.5% BSA (NZYTech, Lisbon, Portugal) was performed for 1 hour. Primary antibodies used were: rabbit anti-aSyn (1:2000, Santa Cruz Biotechnology, Dallas, TX, USA) and anti V5 antibody (1:2000, Life Technologies-Invitrogen, Carlsbad, CA, USA). After overnight incubation with the primary antibody at 4°C, cells were washed with PBS and incubated for 2 hours at RT with Alexa Fluor 488 donkey anti-rabbit (1:2000, Life Technologies-Invitrogen, Carlsbad, CA, USA) and Alexa Fluor 555 donkey antimouse (1:2000, Life Technologies-Invitrogen, Carlsbad, CA, USA). Finally, cells were stained with DAPI (1:5000 in PBS) for 5 minutes and maintained in PBS for epifluorescence microscopy.

Additionally, 48 hours after treatment HEK293 cells stably expressing aSyn-EGFP were fixed with 4% PFA for 20 minutes at RT, followed by a permeabilization step with 0.1% Triton X-100 for 20 minutes at RT. Following permeabilization, a blocking step with 1.5% BSA was performed for 1 hour. Primary antibody used was mouse anti-Syn1 (1:1000, BD Biosciences, Heidelberg, Germany). After overnight incubation with the primary antibody at 4°C, cells were washed with PBS and incubated for 2 hours at RT with Alexa Fluor 555 donkey anti-mouse (1:1000, Life Technologies-Invitrogen). Cells were finally stained with DAPI (1:5000 in PBS) for 5 minutes and mounted in coverslips for epifluorescence microscopy.

#### Microscopy

Pictures were taken on a Leica DMI 6000B microscope, with an objective of 40x. The camera used was a Leica DFC 350 FX R2. The filter modes used were Y3 (666 nm), GFP (516 nm) and A4 (451 nm). The software used was Leica Application Suite AF.

#### Determination of the size of aSyn inclusions

Average inclusion size was determined using *ImageJ*. For each image, individual inclusions were identified, the image was then duplicated and a Gaussian blur filter was applied to one of the images with a sigma radius value of 10. The blurred image was subtracted from the original and the image threshold of the image obtained was adjusted. The resulting image was used to analyse the particle size directly with the analyse particles plug-in.

#### Quantification of aSyn inclusions

H4 cells transfected with aSyn were scored based on the aSyn inclusion pattern and classified into four different groups: cells without inclusions, cells with less than six inclusions, cells between six and ten inclusions and cells with more than ten inclusions. Results were expressed as percentage of the total number of transfected cells obtained from three independent experiments. For the PFF-treated cells, the presence or absence of inclusions was assessed and the data were expressed as percentage of cells with inclusions.

### *In vivo* studies

#### Growth and maintenance of *C. elegans* strains

Animals were generally maintained in Petri dishes on solid nematode growth medium (NGM) seeded with OP50 strain of *E. coli* as previously described (Brenner, 1974) and synchronized animals were prepared by the standard bleach method (Stiernagle, 2006). For liquid culture media, after hatching, larval stage L1 animals were incubated at 20°C in liquid S-medium enriched with heat-killed of *E. coli* OP50 strain as a food source as previously described (Couillault & Ewbank, 2002; Stiernagle, 2006).

#### Plasmids constructs and generation of transgenic lines

pNB30 (*dat-1p::gfp*) and pNB50 (*dat-1p::SNCA^Wt^*) vectors expression were produced by a gene synthesis manufacturer (GenScript). For amplification, the vectors were transferred in competent bacteria (One Shot TOP 10, Thermo Fischer Scientific) in LB media overnight at 37°C under agitation at 200 rpm. For microinjection, the total genomic DNA from plasmids was extracted from fresh transformed bacteria using a plasmid DNA Maxi Prep kit (Nucleobond XtraMaxi EF, Macherey-Nagel). Transgenic control GFP fluorescent marker expressing line (NB30; *nbIs50[dat-1p::gfp]*; named *control^GFP^* in this study) and double transgenic line coexpressing the full length wild-type human gene of aSyn (417 bp) and the GFP proteins (NB50; *nbIs70[dat-1p::gfp;dat-1p::SNCA^Wt^]*; named *aSyn* in this study) were generated by microinjection following standard transformation techniques (Mello & Fire, 1995). For this, we injected in the gonads of the animals (i) for the control^GFP^ line, GFP fluorescent marker plasmid (NB30) at the final concentration 40 ng/μl and (ii) for the aSyn line, the *SNCA* gene (NB50) in association with the GFP plasmid (NB30) at the final concentration of 60 ng/μl and 40 ng/μl respectively. To obtain stable transgenic lines, integration of extrachromosomal arrays were induced by gamma irradiation exposures (42 Gy; CellRad Faxitron, Edimax) of GFP-expressing preselected L4 animals. Lines showing stable and full transmission of the *gfp* transgene expression were cloned and backcrossed three times with the *N2* wild-type referenced background. The strains *N2* and *cat-2 (e1112)* (deficient mutant of ortholog of human tyrosine hydroxylase) (Lints & Emmons, 1999) were provided from the *Caenorhabditis Genetics Center* (University of Minnesota, St. Paul, MN).

#### Drug treatment

Larval stage L1 after hatching were grown in liquid media in presence of doxycycline at the final concentration of 15 μM during 3 days at 20°C under agitation (70 rpm). To avoid small progeny larvae, young adult animals before egging (approximatively 72h of incubation) were put on solid NGM media and were individually picked to put back in culture liquid media with doxycycline treatment until day 6. L-Dopa (*Sigma-Aldrich*, St. Louis, MO, USA) was dissolved in water at 10 mM and young adult stage animals were incubated at a final concentration of 1 mM during 2h hours at 20°C under agitation (70 rpm).

#### Behavioural analysis

To evaluate dopaminergic behaviours induced by aSyn we used a method described by Vidal-Gadea *et al* (Vidal-Gadea et al, 2011). Behavioural test of young adult animals was preceded by an incubation of 2 hour in M9 medium without food source at 20°C without doxycycline in presence or absence of L-Dopa (1 mM). To motivate the movement of the worms, assay plates containing bacteria were prepared by spreading a drop of OP50 *E. coli* in the center of the plates followed by incubation at 37°C overnight. A drop of 5 μl of about 10 worms was placed on the assay plate and the assays began 5 min after the drop was absorbed by the agar, i.e., at a time where the worms were able to escape and start crawling (recordings were done for 5 min on a dry zone). We performed movies during 1 min (7 fps, AZ100M microscope from Nikon) on a total of 30 worms per condition for each experiment. Crawling velocity was analysed by tracking the centroid of animals by video using the wrMTrck plugin of *ImageJ* software (Nussbaum-Krammer et al, 2015). Assays were performed with three independent experiments, n = 30 animals per experimental condition and per experiment.

#### Immunostaining of *C. elegans*

Hermaphrodite worms were synchronized and young adults were prepared using a variation of the *freeze-crack* method from Albertson (Albertson, 1984). Worms were washed by centrifugation in *M9* and placed on a poly-L-Lysine-coated slide (superfrost) previously incubated with a drop of poly-L-Lysine *(Sigma-Adrich)* for 15 min at 60°C. The samples were incubated with 50 μl of primary antibodies diluted in the blocking solution overnight at 4°C. Monoclonal anti-aSyn antibodies used for this study were *Ab6176* (1:200, *Abcam;* recognizes the N-terminal part of the human aSyn protein, red signal in fig. 4B). The tyrosine hydroxylase enzyme (TH) was detected with the antibody #22941 (1:200, Immunostar, blue signal in fig. 4B). After washing, the worms were incubated in a solution containing the secondary anti-mouse (for TH; Alexa Fluor, Red-548, Life Technologies) and anti-rabbit antibody (for aSyn; Alexa Fluor, Red-647, Life Technologies) for 2 hours at RT. Slides were mounted with 8 μl of anti-fade gold (Life Technologies) and stored at 4°C.

#### Quantification of dopaminergic neurons

Quantification of GFP tagged DA neurons was performed in young adult worms raised at 20 °C. Animals fixed in PBS with 4 % PFA were then mounted on slides. The potential neuroprotective effect of doxycycline was evaluated with an epifluorescence microscope (Apotome.2 imaging system from Zeiss) using an x100 magnification lens. Quantifications were performed from three independent experiments, *n* = 25 animals per experimental condition and per experiment. In a second analysis, images sections of PDE DA neurons were acquired, the surface area and the intensity of the GFP of each neuron was quantified using the *3D object counter* plugin of *ImageJ* software (90). Date were obtained from three independent experiments, *n* = 20 animals per experimental condition and per experiment.

#### aSyn distribution in DA neurons in *C. elegans*

To determine the localisation of the aSyn immunosignal, images of DA GFP tagged neurons were captured with an inverted confocal microscope (*Leica TCS SP8*) using an x63 oil objective. For quantification of aSyn aggregates, stacks of 20 images sections of 0.228 μm per neuron were acquired per animal. Analysis of the aSyn immunostaining distribution was assessed using the *3D object counter* plugin of *ImageJ* software. Assays were performed with three independent experiments, *n* = 20 neurons per experimental condition and per experiment.

#### Statistical analyses

All data were obtained from at least three independent experiments and expressed as mean ± SEM or mean ± SD. Comparisons were made using ANOVA followed by *Tukey’s* or *Dunnett’s* post-hoc test and Student’s t test whenever required. Differences were considered as statistically significant at p < 0.05. Statistical analyses were carried out with GraphPad Prism 5 (San Diego, California, USA) and EXCEL (Microsoft, Seattle, WA, USA).

## Acknowledgments

ADM is supported by a postdoctoral fellowship from the Galician Government (Programa de axuda á etapa posdoutoral, XUGA, GAIN, ED481B 2017/053).

The authors want to thank Dr. Filippo Favretto for his help and advice with the DLS measurements, Patrícia Santos for her help with the SDD western blots.

TFO is supported by the Deutsche Forschungsgemeinschaft (DFG, German Research Foundation) under Germany’s Excellence Strategy - EXC 2067/1-390729940, and by SFB1286 (project B8). VP is the recipient of a Thesis scholarship from Association France Parkinson. PPM is supported by the Innovative Medicines Initiative 2 Joint Undertaking under grant agreement No 821522 (PD-MitoQUANT; this Joint Undertaking receives support from the European Union’s Horizon 2020 research and innovation program and EFPIA and Parkinson’s UK). This work also benefited from support by Investissements d’Avenir (ANR-10-IAIHU-06) and the Translational Research Infrastructure for Biotherapies in Neurosciences (ANR-11-INBS-0011-NeurATRIS). This work also was supported by PICT 2015 N° 3201. The study was supported by the São Paulo State Foundation for the Support of Research (FAPESP, Brazil; Grant 2014/25029-4). EDB is a recipient of grants from the National Council for Scientific and Technological Development (CNPq, Brazil), CAPES, and the French Committee for the Evaluation of Academic and Scientific Cooperation with Brazil, Grant #848/15). EDB is a CNPq senior research fellow.

## Author Contributions

EDB, RC, RRV, PPM, NB and TFO designed research. ADM, VP, EV and FGL performed research. AK and DFL contributed new reagents or analytic tools. ADM, VP, EF, FGL, AL, SH, EDB, RC, RRV, PPM, NB and TFO analyzed data. ADM, VP, EF, FGL, AL, SH, EDB, RC, RRV, PPM, NB and TFO wrote the paper.

## The paper explained

### Problem

Parkinson’s disease (PD), a devastating and incurable neurodegenerative disease, belongs to a broader family of diseases known as synucleinopathies, known for the accumulation of aggregated of α-synuclein (aSyn). Clinically, these diseases can be readily distinguished, but the molecular underpinnings are still obscure. Therefore, development of new therapeutic strategies against these diseases is of utmost importance due to the burden they cause to patients, their families, and societies as a whole. In this regard, the repurposing of existing drugs emerges as a promising strategy against synucleinopathies.

### Results

In our study, we assessed the effect of the antibiotic doxycycline on aSyn in several cell free, *in vitro*, and *in vivo* models, and obtained strong evidence on the effectiveness of doxycycline against synucleinopathies. Our results show that doxycycline reduces aSyn aggregation, normalizes the levels of reactive oxygen species generated by mitochondria in cells and, in a newly developed *C. elegans* model, alters the subcellular distribution of aSyn in dopaminergic neurons and restores motor alterations.

### Impact

In summary, our results demonstrate the potential of doxycycline in synucleinopathies in different in vitro and in vivo models. We also provide insight into the mechanisms by which doxycycline may inhibit the aggregation of aSyn and into its viability as a possible treatment for synucleinopathies.

## Expanded view figures

**Supplementary figure 1.**
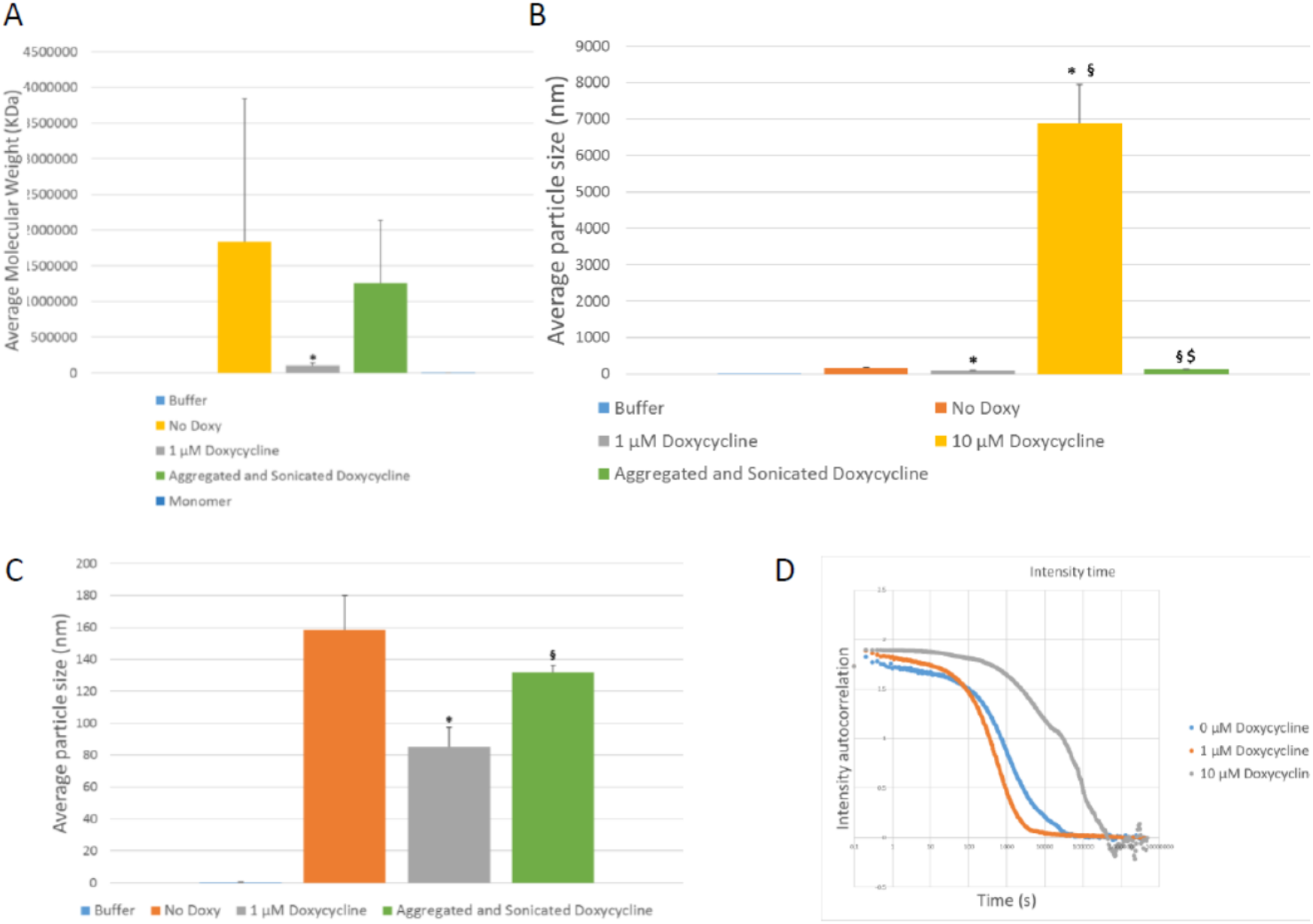
The presence of doxycycline leads to changes in average molecular weight and average particle size of RT-QuiC products. **A.** Average molecular weight of the lower weight species measured by DLS. There are no statistically significant differences amongst groups. B. Average particle size of the RT-QuiC products measured by DLS. Treatment with 10 μM Doxycycline leads to a significantly higher average particle size. **C.** Average particle size of the lower weight species measured by DLS. The presence of 1 μM doxycycline during RT-QuiC leads to a statistically significative decrease in average particle size in comparison with untreated RT-QuiC products before and after a posterior sonication. **D.** DLS measurements of the samples obtained in the RT-QuiC. Above equimolar concentrations higher molecular weight species lead to higher times for the decrease of the intensity correlation. N = 3 in all experiments *p < 0.05 in comparison with 0 μM Doxycycline. §p< 0.05 in comparison with 1 μM Doxycycline. $p< 0.05 in comparison with 10 μM Doxycycline.

**Supplementary figure 2.**
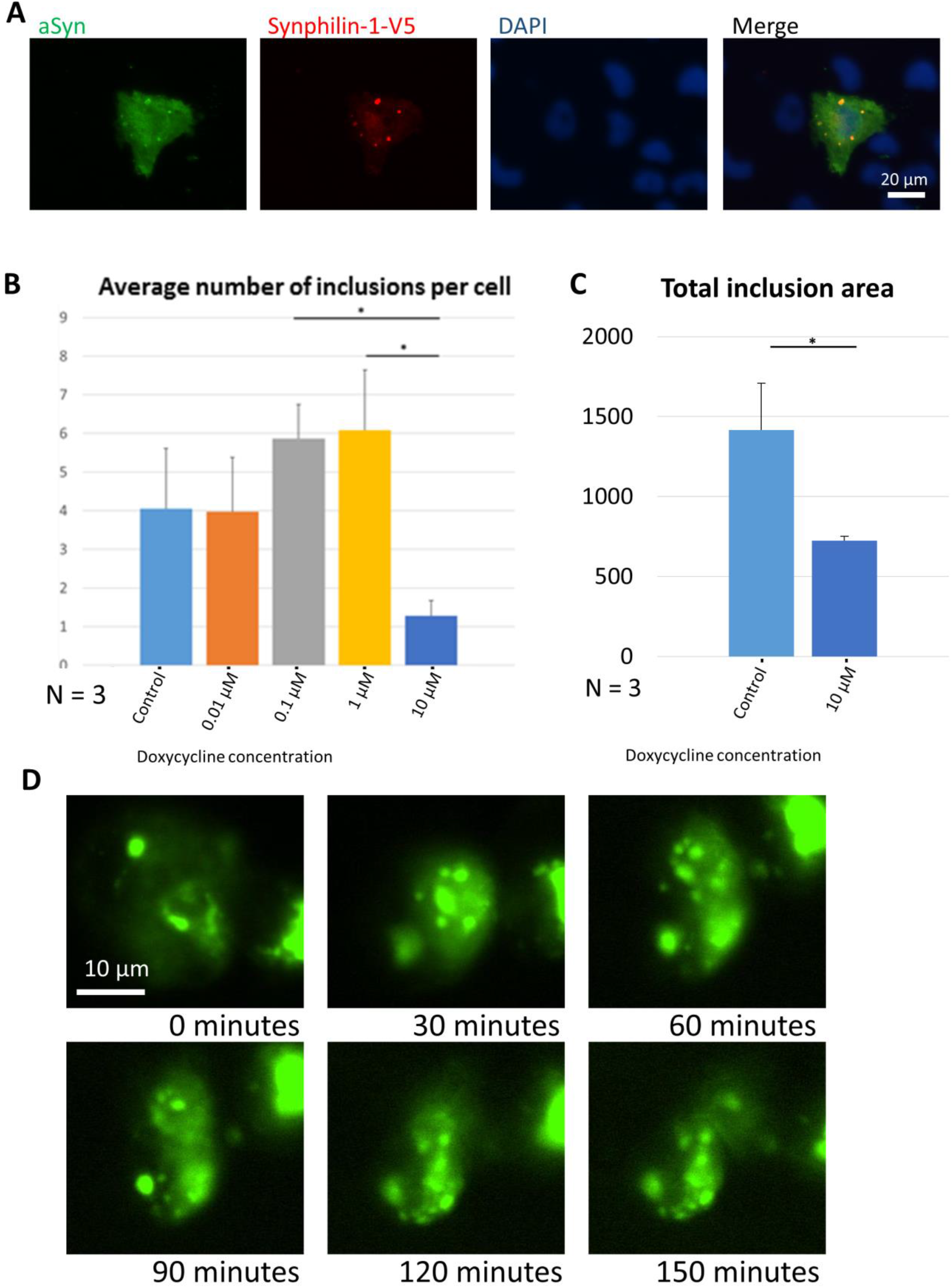
Treatment with doxycycline leads to an increase in the average number of inclusions per cell. **A.** Immunostaining for aSyn and Synphilin-1-V5 shows presence of both proteins in inclusions. **B**. Average number of inclusions per cell. Treatment with different concentrations of doxycycline leads to a significantly different average number of inclusions per cell. Data presented as mean ± SD. N = 3 in all experiments * p < 0.05 **C.** Total inclusion area (in μm^2^) of control cells and cells treated with 10 μM doxycycline. Treatment with 10 μM of doxycycline lead to a significantly different overall inclusion area in cells. Data presented as mean ± SD. N = 3 in all experiments * p < 0.05. **D.** Images of a cell transfected with VNSynph/aSynVC and treated with 10 μM doxycycline at different time points. An apparent increase in the number of inclusions accompanied by a decrease in their size can be seen.

